# From colonisation to chronicity: adaptation of *Mycobacterium abscessus* in the cystic fibrosis lung environment

**DOI:** 10.1101/2025.08.27.672625

**Authors:** Niamh Duggan, Deirdre Keating, Joanna Drabinska, Ciarán J. Carey, Guerrino Macori, Kirsten Schaffer, Siobhán McClean

## Abstract

Chronic infection by opportunistic pathogens is a major contributor to mortality in people with cystic fibrosis (CF). These infections are caused by antimicrobial resistant (AMR) pathogens such as the emerging pathogen, *Mycobacterium abscessus*, a nontuberculous mycobacteria (NTM) which causes recalcitrant infections with high resistance to antibiotics.

*M. abscessus* adapts over time of colonisation to the conditions in the CF lung, hampering effective treatment. The mechanisms underlying this pathoadaptation are poorly understood and are critical for the development of future therapies. Sequential isolate pairs of *M. abscessus* from three people with CF were examined for adaptive changes over time of infection. Genomic analysis confirmed that these isolate pairs were clonal. The late infection isolates showed increased host cell attachment to CF bronchial epithelial cells and increased intracellular survival in macrophages, indicative of adaptation to the CF lung environment. Late isolates also showed changes in their proteomes, including changes in abundance of proteins with roles in intracellular survival and antibiotic resistance. Overall, it is clear that *M. abscessus* can adapt to the CF lung environment and improve its ability to interact with host cells.

**Impact Statement:** Chronic infection by antimicrobial resistant bacteria impacts both the quality of life and mortality in people with CF. We explored the process of adaptation in the emerging pathogen, *Mycobacterium abscessus,* over the course of a chronic infection in the CF lung. While this process has been well-documented in other opportunistic pathogens which colonise the CF lung, limited data exist on this process in *M. abscessus*. Phenotype and proteome changes were assessed in sequential longitudinal clinical isolates of *M. abscessus* obtained from Saint Vincent’s University Hospital (SVUH), Dublin, Ireland. Significant changes in the interactions with human cells were observed in late infection isolates after as little as 33 days. Understanding these processes may reveal new avenues for clinical exploitation and this study reveals some novel adaptations which could be exploited to aid current therapies.

## Introduction

Chronic progressive lung disease due to infections by opportunistic pathogens is the largest contributor to morbidity and mortality in people with cystic fibrosis (CF) (1). CF is the most prevalent life-limiting autosomal genetic disorder in Caucasians, occurring in between one in 3000 and one in 6000 births in populations of European descent (2–4). Impaired chloride ion transport and build-up of mucus on epithelial surfaces impedes normal mucociliary clearance, facilitating chronic bacterial infections that eventually leads to obstructive lung disease and progressive loss of lung function (5–8). Treatment of these infections is complicated by high levels of antimicrobial resistance (AMR) (9).

*Mycobacterium abscessus* is a member of the nontuberculous mycobacteria (NTM) group (10) and an emerging pathogen of concern for people with CF or other respiratory conditions such as chronic obstructive pulmonary disorder (COPD) (11). Pulmonary NTM disease leads to progressive inflammatory lung damage and, notably, can involve prolonged asymptomatic incubation within the lungs, complicating detection and diagnosis (12, 13). The annual incidence of NTMs rose from 3.13 to 4.73 per 100,000 individuals in the US between 2008 and 2015, while the prevalence increased from 6.78 to 11.70 per 100,000 (14). In people with CF, NTM pulmonary infections increased by 3.5% annually between 2010 and 2019 (14). The *M. abscessus* complex encompasses three subspecies*: M. abscessus abscessus, M. abscessus bolletii* and *M. massiliense* (15, 16)*. M. abscessus* is particularly difficult to treat due to its intrinsic and acquired resistance to many classes of antibiotics including macrolides, aminoglycosides, rifamycins, tetracyclines and β-lactams (17). *M. abscessus* pulmonary disease is considered very difficult to eradicate (11), with treatment success rates as low as 45.6% (18).

*M. abscessus* is a rapidly growing mycobacteria and can occur as either a rough or smooth colony morphotype (19). Smooth morphotypes are motile, biofilm-forming and non-cord-forming as they produce glycopeptidolipid (GPL) in their cell walls while rough morphotypes are non-motile, do not form biofilms, and form cords (19, 20). Although *M. abscessus* is an environmental bacterium and many of the cases are thought to be due to independent acquisitions from exposure to its natural reservoirs, nosocomial infections have been reported (21–23). Many of the isolates colonising people with CF cluster into dominant circulating clones (DCCs). Phylogenetic analysis of over 2,000 *M. abscessus* isolates identified seven DCCs which have emerged as recently as 1999, with historical population expansions occurring for six of the seven DCCs in the 1960s (24).

Bacterial adaptation is imperative for persistent colonisation of the CF lung. Many CF-associated pathogens display incredible genetic plasticity and adaptability, enabling them to react rapidly to new niches (25, 26). Extreme selection pressures such as antibiotic treatment can result in rapid adaptation via increased mutation rates (27, 28), but adaptation can also occur more gradually via mutations in specific genes that enhance fitness and virulence of bacterial clones. Mutations in another CF-associated pathogen, *Pseudomonas aeruginosa* are positively selected for, and provide a fitness benefit via convergent evolution (29). Moreover, bacterial populations also sustain multiple distinct clonal lineages to facilitate the rapid exploitation of new environmental niches as they become available (27).

Although the process of adaptation has been well studied in other CF-associated pathogens, studies on the adaptation of *M. abscessus* are limited. *M. abscessus* undergoes within-host evolution, accumulating mutations that enhance persistence, immune evasion, and antibiotic resistance (21, 30). Key adaptive traits include mutations in resistance genes such as *erm*(*41*), *rrl*, and *rrs*, shifts from smooth to rough morphotypes associated with loss of glycopeptidolipids, and changes in transcriptional regulators such as *whiB7* and *sigH*; alterations in fatty acid metabolism, biofilm formation, and cell wall composition, supporting survival in immune-challenged environments such as the CF lung (21, 30).

The aim of this work was to gain a better understanding of the adaptation process of *M. abscessus* over the course of a chronic infection in the CF lung. We examined host-cell interactions, antibiotic susceptibility and virulence factors of three pairs of sequential longitudinal clinical isolates of *M. abscessus* obtained from Saint Vincent’s University Hospital (SVUH) for adaptive changes over time of infection that might contribute to chronic infection. We also compared the proteomes of these isolates to identify possible mechanisms of adaptation.

## Methods

### Bacterial Strains

*M. abscessus* isolates were collected for this study from sputum samples of CF patients attending SVUH and sequential isolates were available for three patients (Table 1). In each case, the earliest available isolate and the most recent isolate available at the time of the study were selected (termed ‘early’ or ‘late’ isolates, respectively) (**Fig. 1A**). Isolates were routinely cultured on Middlebrook 7H11 agar plates (Merck Millipore, Burlington, MA, USA) supplemented with 5% glycerol and 10% oleic acid albumin dextrose catalase (OADC) at 37 °C and in Middlebrook 7H9 liquid medium (BD, Wokingham Berkshire, United Kingdom) supplemented with 5% glycerol, 0.05% Tween 80 and 10% albumin dextrose catalase (ADC) at 37 °C with orbital agitation (200 rpm).

**Figure 1.**
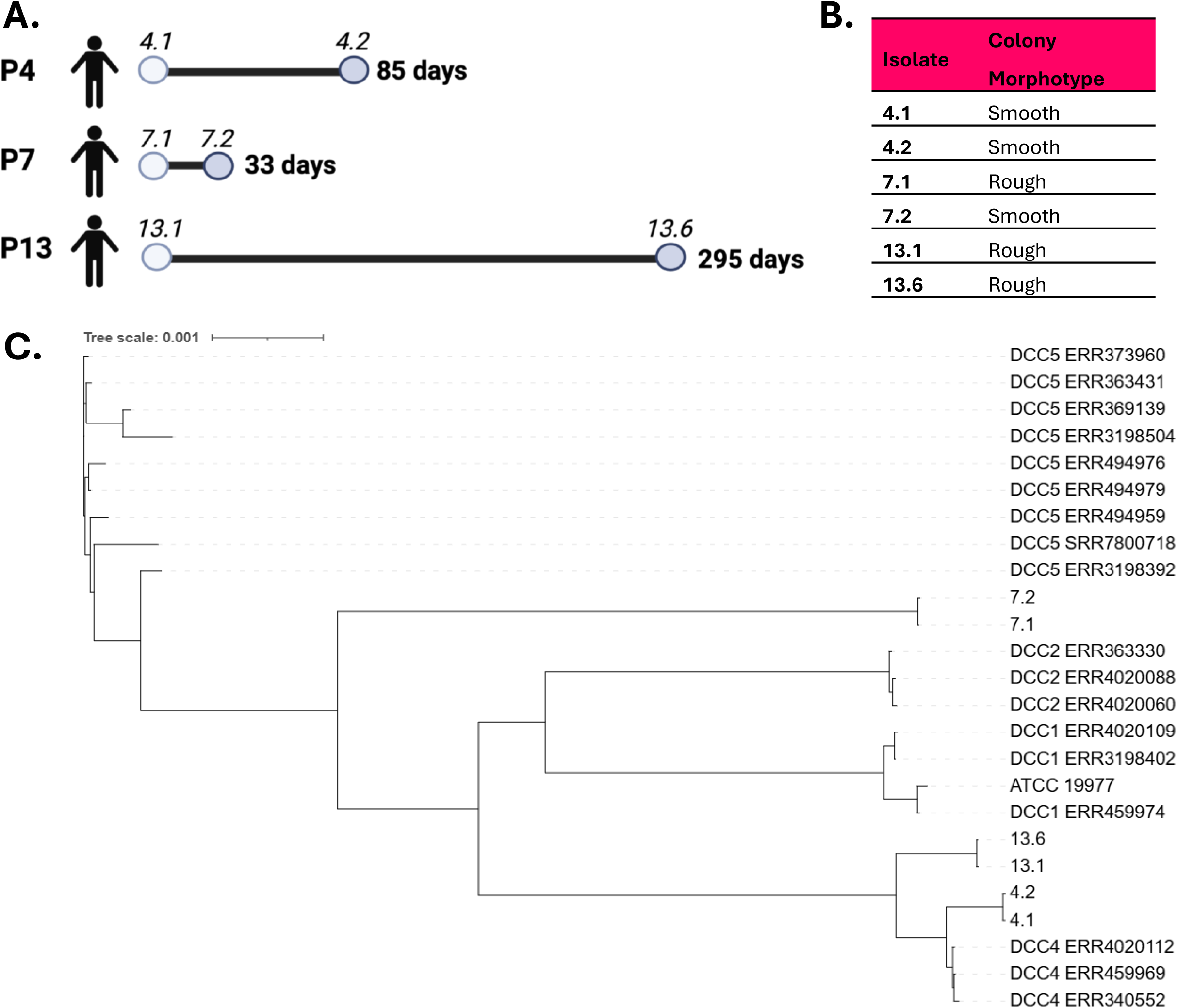
Longitudinal sequential isolates examined. **A)** *M. abscessus* isolates investigated in this study and the timeline of isolation. Timeline shows time from date of first available isolate (early) to date of the last available isolate (late) in the collection. **B)** Colony morphotype of isolates: assessed visually by growth on agar plates after 5 days at 37 °C**. C)** Expanded phylogenetic tree incorporating seventeen isolates from DCC clusters. NovaSeq with 2×150 paired-end reads was used for sequencing and analysis performed on usegalaxy.org (2024). SNPs were determined using Snippy (Galaxy version 4.6.0 +galaxy0). SNP alignment was performed using Snippy Core. The phylogenetic tree was created using Fasttree (34)and iTOL (interactive tree of life) accessed via <u>iTOL: Interactive Tree Of Life</u>.

**Table 1:** Clinical data available on the CF patients during and following the time of *M. abscessus* isolation.

| Patient | Early isolate | Late isolate | Genetics | Class | Modulator therapy | FEV1 Early (Date m/y) | FEV1 Recent (Date) | FEV1 Early | FEV1 Recent | Change (%) |
| --- | --- | --- | --- | --- | --- | --- | --- | --- | --- | --- |
| 4 | 02/02/17 | 28/04/17 | ΔF508/ΔF508 | Class II | Orkambi Oct 2018 – Nov 2019<br>Symkevi & Kalydeco Nov 2019 – Mar 2021,<br>Kaftrio & Kalydeco since Mar 2021 | 08/17 | 03/24 | 2.52L<br>(79%) | 3.13L<br>(97%) | +18% |
| 7 | 30/01/17 | 04/03/17 | ΔF508/ΔF508 | Class II | Orkambi Aug 2017 – Nov 2018<br>Symkevi & Kalydeco Nov 2018 – Oct 2020,<br>Kaftrio & Kalydeco since Oct 2020, | 03/17 | 11/24 | 2.38L<br>(56%) | 2.92L<br>(69%) | +13% |
| 13 | 07/04/22 | 27/01/23 | Heterozygous<br>non ΔF508 | Class I | None | 01/23 | 11/24 | 2.49L<br>(48%) | 1.93L<br>(37%) | -11% |

### DNA extraction and sequencing

*M. abscessus* isolates were inoculated into Middlebrook 7H9 liquid medium (BD, Wokingham Berkshire, United Kingdom) supplemented with 5 % glycerol, 0.05 % Tween 80 and 10 % albumin dextrose catalase (ADC) at 37 °C with orbital agitation (200 rpm) @37 °C for 96 hours. DNA was extracted with mechanical beads using a MP FastPrep® tissue homogeniser and precipitated using ethanol. Purification was performed with AMPure XP (Beckman coulter) beads. DNA was quantified and checked for purity using Nanodrop (Thermoscientific) and Qubit® (Invitrogen) according to manufacturers’ instructions. The purified *M. abscessus* isolates were sequenced using illumina NovaSeq 2X 150bp (Illumina, San Diego, CA, USA). The resulting fastq files were used for subsequent analysis.

### Core genome phylogeny of isolates

*M. abscessus* paired isolates were sequenced using NovaSeq 6000 genome sequencing platform with 2 x 150 paired end reads. Analysis of isolates was performed using bioinformatic tools available on usegalaxy.org (31). Raw data (fastq) was first assessed for quality using fastqc (Galaxy Version 0.74+galaxy1). Pre-processing was performed with fastp (Galaxy Version 1.0.1+galaxy0); adaptors were removed and quality scores set to include only reads with quality scores >Q30. Reads were assembled using SPAdes (Galaxy Version 1.1.0+galaxy2) and the resulting genomes were annotated using Prokka (Galaxy Version 1.14.6+galaxy1). Genomes for individual isolates were assembled using Shovill (Galaxy Version 1.1.0+galaxy2). Reads were initially trimmed using Trimmomatic to remove common adapters and then assembled using SPAdes (https://github.com/tseemann/shovill?tab=readme-ov-file#shovill). Single Nucleotide Poly-morphism (SNP) identification and analysis was subsequently performed on the assembled genomes using kSNP3.0)(32).The resulting parsimony output was visualised using iTOL v6. A phylogenetic tree was compiled using SNP analysis from patient samples, published known clonal complex genomes from DCC1, 2, 4 and 5 and *M. abscessus abscessus* ATCC 19977 (selected from (24)). SNPs present between the isolates and *M. abscessus abscessus* reference genome ATCC19977 were determined using Snippy (Galaxy version 4.6.0 +galaxy0) (33). Multiple SNP alignments created were combined using Snippy-core and the resulting output was used to create a newick (.nhx) file on Fasttree (Galaxy Version 2.1.10+galaxy1) (34). This was visualised using iTOL (interative tree of life) iTOL: Interactive Tree Of Life.

### Screening for genetic resistance

Assembled genomes were uploaded to CARD RGI (Antibiotic Resistance Database resistance gene identifier) which provides data, models, and algorithms relating to the molecular basis of antimicrobial resistance. CARD reference 10.1093/nar/gkac920

### Biofilm formation

Starter cultures were diluted to OD_600nm_ 0.1 and 200 μL added to wells of round bottom 96-well plates in Middlebrook 7H9 broth supplemented as described. Plates were wrapped in parafilm and tinfoil to prevent evaporation and incubated for 7 days at 37 °C and 5% CO_2._ The OD_600nm_ was measured using the Biotek Syngery H1 microplate reader (Mason Technology, Dublin, Ireland). The media containing planktonic bacteria was removed and biofilms fixed by the addition of 150 μL of methanol for 5 mins. Wells were washed three times with 150 μL of phosphate buffered saline (PBS) (Sigma-Aldrich, St. Louis, MO, USA) and stained with crystal violet (0.1%, 150 μL) at room temperature for 30 mins. The crystal violet was removed and the plate washed with water and then with PBS before drying for 30 mins at room temperature. The crystal violet was then solubilised with 200 μL of 96% ethanol for 30 mins and mixed. An aliquot from each well (150μL) was added to a fresh well to read the OD_590nm_ on the Biotek Syngery H1 microplate reader. The OD_590nm_/OD_600nm_ ratio was then calculated to assess biofilm production.

### Sliding motility on 0.3% agar

Sliding motility was assessed in 0.3% agar plates prepared (Sigma-Aldrich, St. Louis, MO, USA) with Middlebrook 7H9 media (BD, Wokingham Berkshire, United Kingdom) and dried overnight. Mid-log (OD_600_) bacterial cultures (10 μL) were dropped onto the centre of the plates which were incubated upright at 37°C and 5% CO_2_ for 5 days. Each isolate was examined in duplicate and two perpendicular diameters were measured per plate. Plates images were also captured using the Vilbur E-box imaging system.

### Assessment of antibiotic susceptibility

Antibiotic susceptibility was measured on lawns of each isolate prepared by adjusting mid-log cultures to a McFarland standard of 0.1 and plating 350 μL aliquots on Mueller Hinton 145 mm agar plates (Neogen, Lansing. MI, USA) which were spread using a cotton swab in three directions. Antibiotic disks impregnated with azithromycin (AZM), kanamycin (K), ciprofloxacin, (CIP), cefoxitin (FOX), rifampicin (RID) or amikacin (AK) (Oxoid, Hampshire, UK) were then placed on the agar using a sterile tweezers and plates were incubated for 5 days at 37°C and 5% CO_2_. Zones of inhibition were measured in at least two directions after 5 days growth and the mean was used for statistical analysis. Each experiment was repeated three times.

### Determination of bacterial attachment to human CF epithelial cells

The CF human bronchial epithelial cell line, CFBE41o-(35, 36) was routinely cultured in T25 and T75 flasks precoated with a coating buffer composed of minimal essential media (MEM) containing 10% w/v bovine serum albumin (BSA), 1% v/v collagen (Sigma Aldrich) and 1% v/v fibronectin (Sigma Aldrich) and incubated at 37°C in a humidified incubator at 5% CO_2_. To quantify bacterial attachment, CFBE41o^-^ cells were seeded on fibronectin coated 24-well plates at a density of 4 x 10^5^ cells / well in antibiotic free medium and incubated at 37°C, 5% CO_2_ overnight. Bacteria were cultured to a mid-log OD_600_ of ∼ 0.6 and then homogenized through a 30 G needle 30 times to disperse bacteria before dilution to a final multiplicity of infection (MOI) of 5:1 in MEM without antibiotic. Dilutions were then plated to accurately assess the CFU/mL. MEM (500 μL) and bacterial culture (500 μL) were added to each well and the plates centrifuged at 252 x *g* for five minutes and incubated for 30 minutes at 37°C, 5% CO_2_ to facilitate bacterial adherence. Unattached bacteria were removed by aspiration and the wells were washed three times with PBS. The cells were lysed with 500 μL of 0.25% Triton X-100 in PBS for 20 mins at room temperature (RT), lysates removed by scraping with a pipette tip and then serially diluted in Ringer’s solution. Each dilution (100 μL) was spread onto Middlebrook 7H11 plates supplemented as described above in duplicate and incubated at 37°C for 5 days at which time the resulting colonies were counted.

### Measurement of bacterial uptake and survival in U937 cells

The human monocytic U937 cell line (37) was maintained as a suspension culture in RPMI-1640 supplemented with L-glutamine (2mM), sodium bicarbonate (2g/L), 1% v/v sodium pyruvate, 1% v/v HEPES buffer, 1% v/v streptomycin/penicillin, 10% v/v FBS and 5 g/L D-glucose (Sigma-Aldrich, St. Louis, MO, USA). U937 cells (5 x 10^5^ cells / mL) were differentiated in 24-well plates by the addition of 15 ng/mL of phorbol 12-myristate 13-acetate (PMA) for 48h in full RPMI-1640 medium. Cells were washed in PBS and incubated in RPMI-1640 medium for up to 24h before inoculation. Bacteria were cultured to a mid-log OD_600_ of ∼ 0.6 and dispersed and diluted to 2.5 x 10^6^ CFU / mL in RPMI-1640 medium. U937 cells were washed twice with PBS, and 1 mL of bacterial culture (MOI 5:1) added to duplicate wells. Plates were centrifuged at 1100 x *g* for 5 min and incubated at 37°C, 5% CO_2_ for 2h to allow for uptake. The inocula were serially diluted in Ringer’s solution and plated in duplicate to confirm bacterial cell numbers added. After 2h, extracellular bacteria were removed, and the cells washed twice with PBS, and 1 mL of RPMI-1640 medium containing 1 mg/mL amikacin was added to each well and incubated for 2h at 37°C, 5% CO_2_ to kill any remaining extracellular bacteria. The wells were then washed five times with sterile PBS before cells were lysed with 0.25% v/v Triton X-100 for 15 minutes at RT. The final wash was plated on Middlebrook 7H11 agar to confirm sterility. The wells were scraped with a pipette tip and cell lysates were serially diluted in Ringer’s solution before plating on Middlebrook 7H11 agar plates in duplicate and incubating at 37°C for 5 days for CFU enumeration. Intracellular survival was determined in 24-well plates using the same procedure, with additional steps. After bacterial uptake, wells were washed once with PBS, and fresh media with 1 mg/mL amikacin was added to each well to kill any extracellular bacteria before incubation at 37°C, 5% CO_2_ for up to 7 days. Wells were scraped and lysed for plating on Days 1, 3, 5 and 7 in duplicate.

### Whole proteome analysis

Isolates were cultured for three days in Middlebrook 7H9 media (10 mL) and then pelleted by centrifugation at 4000 x g for 10 mins. Pellets were resuspended in 2 mL of buffer comprised of 50 mM ammonium bicarbonate, 10 mM magnesium chloride, 0.1% sodium azide, 1mM EGTA, 7M urea, 2M thiourea and 1x cOmplete^TM^, EDTA-free protease inhibitor cocktail (Roche, Basel, Switzerland). Glass Zirconia/silica (0.1mm) beads (Fisher Scientific, Hampton, New Hampshire, United States) were added at a ratio of 1:3 (v/v) and lysed in a FastPrep-24 Bead Beater (MP Biomedicals, Santa Ana, CA, USA) for 1 min 30s at 5.0 m/s. Lysates were centrifuged at 17,000 x g for 30 mins at 4°C to remove debris and any remaining intact cells. Supernatants were precipitated overnight in 25% trichloroacetic acid (TCA) at 4°C. Precipitates were centrifuged at 17,000 g for 15 mins at 4 °C, before washing twice in 80% acetone and resuspending in 50 mM ammonium bicarbonate and 1M urea. Samples were centrifuged at 10,000 g at 4 °C for 30 mins before resuspension in 2mL 8M urea buffer, 50mM Tris-HCl pH 8. The protein concentration post lysis was determined with a Nanodrop spectrophotometer DS-11+ (DeNovix, Wilmington, DE, USA). Dithiothreitol (1M) was added (10 μL/mL lysate) and incubated at 56°C for 30 mins before addition of iodoacetamide (1M) (55 μL/mL lysate) and incubation at RT in the dark for 20 mins. The iodoacetamide was quenched with dithiothreitol (final concentration of 20mM) and the protein diluted to 1mg/mL. Samples were dialysed in pre-soaked Snakeskin tubing (cut-off of 3.5 kDa, Thermo Fisher, Waltham, MA USA) in 100 mM ammonium bicarbonate overnight at 4 °C at 121 rpm, with a further incubation of at least 4 h in fresh ammonium bicarbonate. Trypsin digested proteins were added to fresh tubes and dried at 65 °C and resuspended in 20 μL 0.5% Trifluoroacetic Acid (TFA) (v/v) in MilliQ water before being purified using ZipTips (Merck Millipore, Burlington, MA USA) as per manufacturer’s instructions and measure protein concentrations determined as before.

The samples were analysed on a Bruker TimsTOF Pro mass spectrometer connected to a Evosep One chromatography system. Peptides were separated on an 8 cm analytical C18 column (Evosep, 3 μm beads, 100 μm ID) using the pre-set 33 samples per day gradient on the Evosep one. A trapped ion mobility (TIMS) analyser was synchronized with a quadrupole mass filter to enable the highly efficient PASEF (Parallel Accumulation Serial Fragmentation acquisition) procedure with acquisition rates of 100 Hz. The accumulation and ramp times for the TIMS were both set to 100 ms, with an ion mobility (1/k0) range from 0.62 to 1.46 Vs/cm. Spectra were recorded in the mass range from 100 to 1,700 m/z. The precursor (MS) Intensity Threshold was set to 2,500 and the precursor Target Intensity set to 20,000. Each PASEF cycle consisted of one MS ramp for precursor detection followed by 10 PASEF MS/MS ramps, with a total 658 cycle time of 1.16 s. Protein identification and label[free quantitative (LFQ) analysis were conducted using MaxQuant (version 1.2.2.5, https://maxquant.org/) by searching against the reference *M. abscessus* strain ATCC 19977 (561007). The average of LFQ intensities per sample type were calculated and statistical analyses performed in Excel. A Log2 Fold Change cutoff of ≥1.5 was used to determine significant changes in abundance per condition when comparing early and late isolate samples (t-test *p*[≤[0.05). In order to gain a better understanding of the functions of these proteins, individual Blast searches were performed for each against the *M. tuberculosis* database H37RV (https://www.uniprot.org/uniprotkb/O06619/entry). Assignments of function were then completed using the Tuberculist server developed by the Institut Pasteur (http://genolist.pasteur.fr/TubercuList) (38).

## Results

### Sequential isolate pairs confirmed as clonal

The three isolate pairs (P4, P7, and P13) representing different time frames (**Fig. 1A)** from the earliest available isolate to the most recent isolate were selected (termed the early and late isolate respectively each case). P4 isolates both exhibited a smooth morphotype, while both P13 isolates showed a rough morphotype (**Fig. 1B**). The only pair that showed a change in morphotype was that from Patient 7, where the early isolate was rough and the late isolate was smooth, in contrast to the usual change with time.

Whole genome sequencing confirmed each isolate pair was clonal via analysis of core insertion/deletion/SNPs. All isolates were identified as *M. abscessus* subsp. *abscessus* with 99% certainty by Multilocus sequence typing (MLST) (**Table 2**). P4 isolates and P13 isolates cluster with DCC 4 while P7 isolates did not cluster with any of the selected DCC isolates. The isolates were further mapped to three isolates from each of the *M. abscessus* DCC clusters DCC1, DCC2, DCC4 and DCC5 (**Supplementary Table S1**) (24). This was subsequently expanded to include an extra seven strains from DCC5, as this appeared to be the closest match to the unmapped P7 isolates. P4 and P13 isolates were confirmed to cluster with DCC4 (**Fig. 1C**). Interestingly, the isolates included in this DCC are all CF patient isolates and two have an Irish source, including one from SVUH, which may suggest these isolates have a common ancestor or are genetically linked. P4 isolates were closely linked to two of the reference isolates, DCC4 ERR4020112 and DCC4 ERR459969, both of which were Irish in origin. P7 isolates did not cluster with any of the DCC groups included in this analysis (**Fig. 1C**). This may suggest it is perhaps an independent environmental acquisition or that the inclusion of more points of reference are needed to establish its origin.

**Table 2:** Genus, sequence type and subspecies of *M. abscessus* identified by whole genome sequencing.

| Isolate | Sequence type | Clonal complex | Core Insertion/<br>Deletion/SNPs | % *M. abscessus*<br>Matched | Other organism<br>matched | Subspecies<br>based on<br>MLST |
| --- | --- | --- | --- | --- | --- | --- |
| 4.1 | 101 | 4 |  | 100 | N/A | *abscessus* |
| 4.2 | 101 | 4 | 0 | 100 | N/A | *abscessus* |
| 7.1 | 129 | Not assigned |  | 100 | N/A | *abscessus* |
| 7.2 | 129 | Not assigned | 0 | 100 | N/A | *abscessus* |
| 13.1 | 97 | 4 |  | 99 | 1% carnobacterium | *abscessus* |
| 13.6 | 97 | 4 | 13 | 100 | N/A | *abscessus* |

### No changes in biofilm production or sliding motility in late isolates

Biofilm production in *M. abscessus* in pulmonary infections in CF aids the establishment of chronic infection (39–41). *M. abscessus* has been reported to become more resistant to antibiotics within biofilms, showing the development of resistance to cefoxitin, amikacin and clarithromycin *in vitro* (42, 43). However, none of the sequential isolates examined in this study formed strong biofilms and there was no change between early and late isolates (**Fig. 2A**).

**Figure 2.**
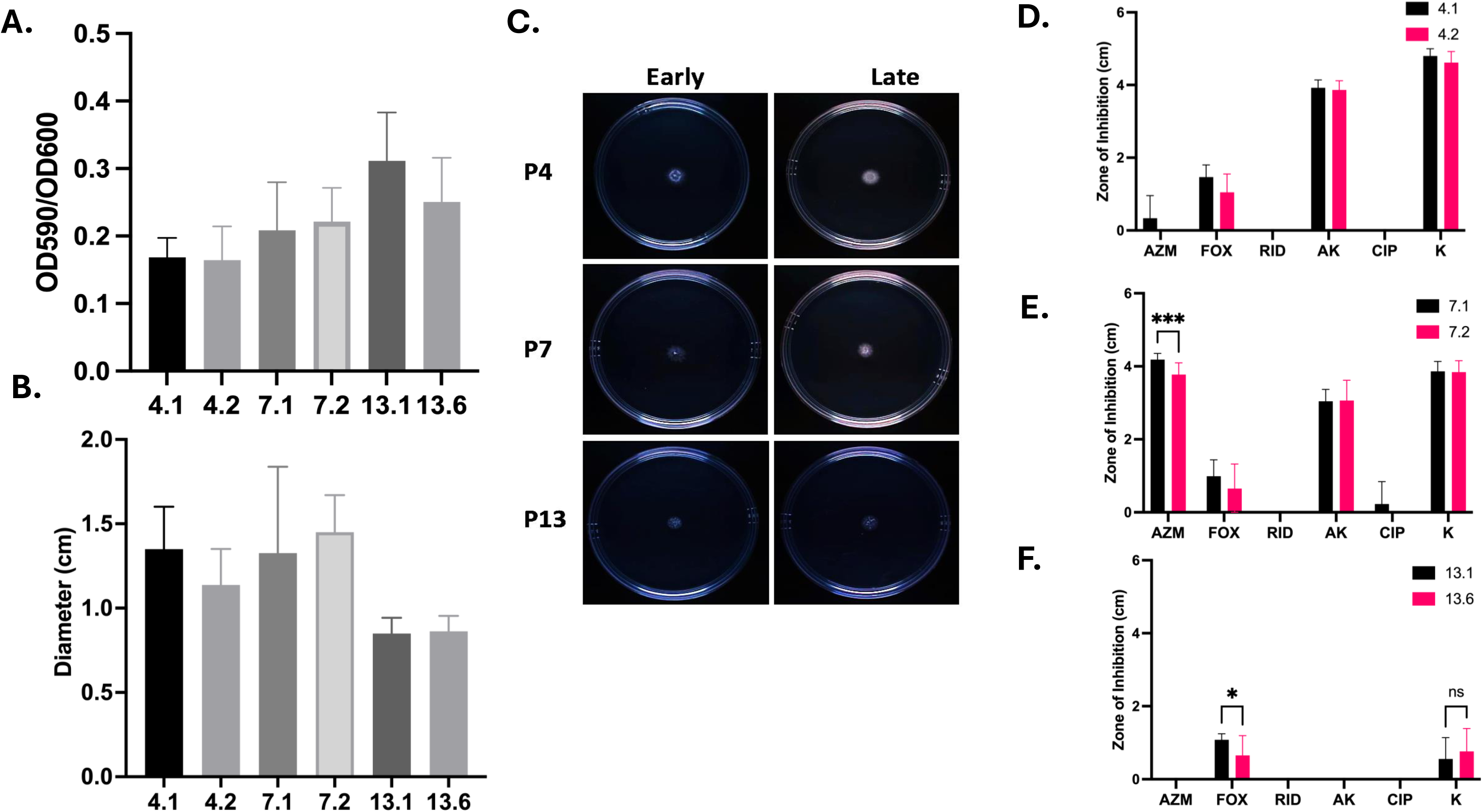
Phenotype adaptations in late infection isolates. **A.** Assessment of biofilm production in P4, P7 and P13 early and late isolates as determined by crystal violet staining. Data represent the mean of two independent experiments, each with five replicate wells per isolate. Data were analysed by unpaired non-parametric Mann Whitney tests. **B.** Assessment of sliding motility in P4, P7 and P13 isolates as measured by growth on 0.3% agar. Data represent mean of two independent experiments, each with three replicate plates per isolate, analysed by unpaired non-parametric Mann Whitney tests. **C.** Representative images of motility plates to assess sliding motility measured on 0.3% agar. Images are representative of two independent experiments with three replicate plates per isolate in each experiment. **D,E,F)** Antibiotic susceptibility of *M. abscessus* clinical isolates from P4 (D), P7 (E) and P13 (F) respectively tested using antibiotic discs impregnated with the following antibiotics: Azithromycin (AZM); cefoxitin (FOX); rifampicin (RID); amikacin (AK); ciprofloxacin (CIP) and kanamycin (K). Data shown are the mean of four independent experiments. Data were analysed by multiple unpaired non-parametric Mann Whitney tests.

*M. abscessus* exhibits passive movement on semi-liquid surfaces known as sliding motility which is caused by bacteria dividing and pushing outward from a point of origin through releasing surfactants (44, 45). Sliding motility was assessed because of its role in colonisation of the host lung surface (46). Similar to biofilm production, there was no dramatic difference in motility between early and late isolates (**Fig. 2B**). All isolates showed minimal movement outward from the point of origin (**Fig. 2C**).

### Late isolates did not show dramatic changes in antibiotic susceptibility

CF isolates are routinely exposed to antibiotics in the CF lung. As expected, analysis of the sequential isolate genomes highlighted that resistance genes associated were detected in all 6 clinical samples (Table S2). There were no differences in the presence of resistance genes in paired isolates from P4 or P7, while P13 isolates showed an increase in SNPs in genes associated with resistance to aminoglycosides between early and late isolates (Supplemental table (S2)).

The isolates were then evaluated for their susceptibility to common classes of antibiotics used clinically. The P4 and P7 isolates showed similar antibiotic susceptibility profiles, with sensitivities to amikacin and kanamycin, but low to no sensitivity to cefoxitin (**Fig. 2D**). The early P4 isolate showed some growth inhibition in response to azithromycin, while the late isolate was fully resistant. Similarly, there was a significant reduction in sensitivity to azithromycin in the late P7 isolate (p ≤ 0.001) (**Fig. 2E**). Neither early nor late P4 or P7 isolates showed sensitivity to rifampicin or ciprofloxacin. Isolates from Patient 13 (**Fig. 2F**) showed little susceptibility to any of the antibiotics tested. There was a small zone of inhibition by cefoxitin in the early isolate which was significantly reduced in the late isolate (p ≤ 0.05), and low levels of response to kanamycin, however both isolates were resistant to all other classes of antibiotics tested.

### Late infection isolates show increased host cell attachment

The ability to attach and adhere to epithelial cells within the CF lung is critical for successful colonisation and survival of *M. abscessus* (47). Previous work in our lab demonstrated increased epithelial cell attachment in sequential *B. cenocepacia* CF isolates over time of colonisation (48). Late infection isolates from P4 and P7 showed significantly increased attachment to CFBE41o-cells *in vitro* relative to their early counterparts (**Fig 3A** and **Fig 3B**). The early P13 isolate showed greater attachment to CFBE41o-cells than the P4 and P7 isolates, at 5% attachment which appeared to be further enhanced in the late P13 isolate, although it was not statistically significant (**Fig 3C)**.

**Figure 3.**
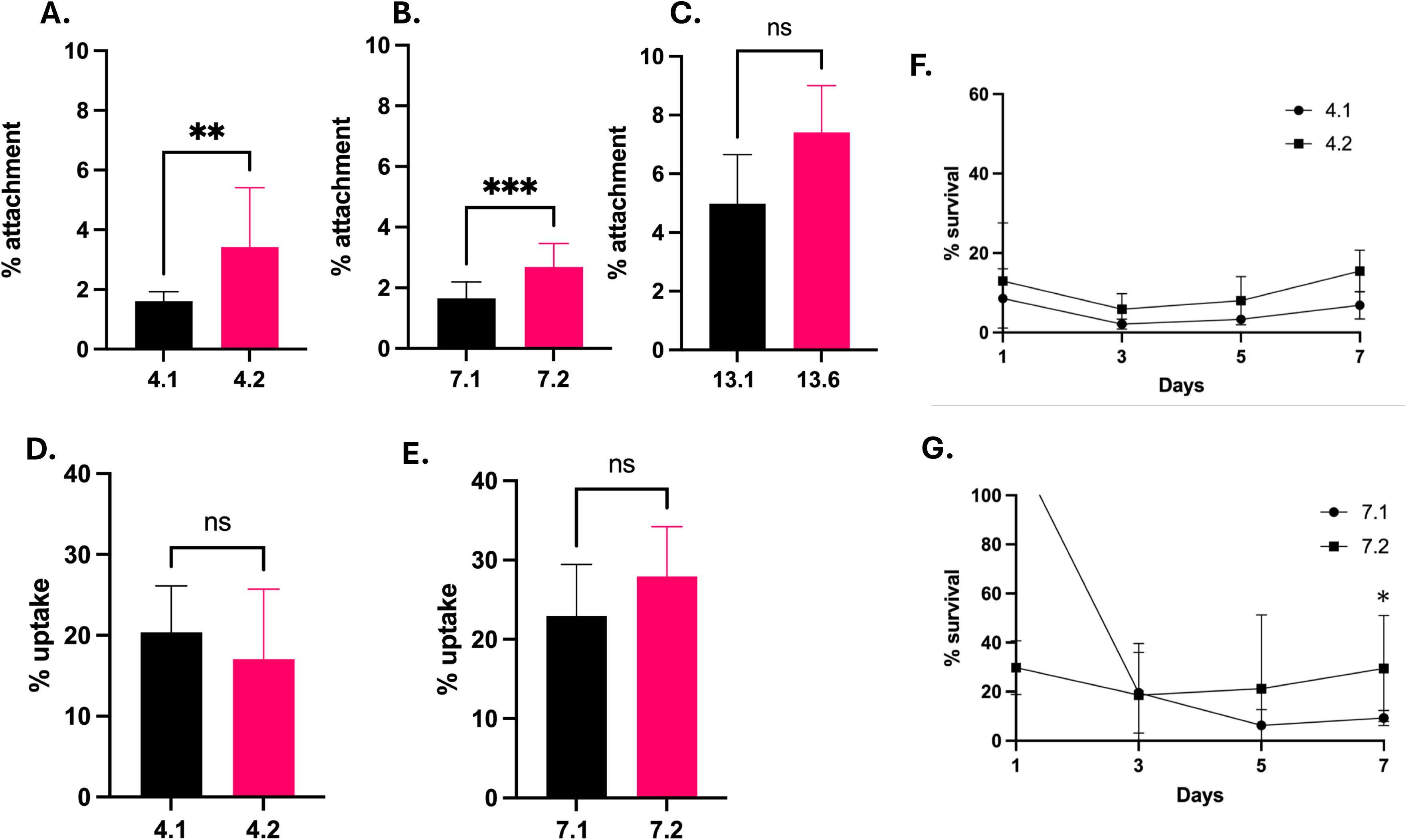
Changes in host-cell interactions. A, B, **C** Attachment of early and late clinical isolates from P4 (A), P7 (B) and P13 (C) to CFBE41o-cells. Results represent the mean of three independent experiments performed in duplicate. Statistical analysis was performed by unpaired non-parametric Mann Whitney tests. **D & E.** Uptake of isolates from P4 and P7 by U937 macrophages over 2 h. Data shown represents the mean of three independent experiments performed in duplicate. Statistical analysis was performed by unpaired non-parametric Mann Whitney tests**. F & G.** Intracellular survival of P4 and P7 isolates over 7 days in U937 macrophages relative to uptake. Data shown represent the mean of three independent experiments performed in duplicate. Statistical analysis was performed by unpaired non-parametric Mann Whitney test on individual days.

### Late infection isolates demonstrate increased intramacrophage survival

The ability to survive intracellularly in phagocytic cells is a hallmark of *M. abscessus* infection and is critical for its persistence in the host (49, 50). Increased expression of universal stress proteins and secretion systems are thought to be key to their survival inside the macrophage, leading to subsequent dissemination (51). The uptake of both P4 and P7 isolates by U937 macrophages was comparable, at approximately 20% uptake (**Fig. 3D** and **Fig. 3E**). Notably, there were no significant differences in macrophage uptake between the early and late isolates in either sequential pair. The capacity to survive intracellularly was slightly higher in the late P4 isolate over 7 days (**Fig. 3F**). By Day 5, there was an increase in cell numbers indicative of intracellular replication inside the macrophage cells, which continued on Day 7, demonstrating an increased ability to survive and replicate inside macrophage cells. A similar trend was observed with the late isolate from P7 (**Fig. 3G**). There was an unexpected high intramacrophage CFU demonstrated by the early P7 on Day 1, perhaps suggesting short-term resistance to oxidative burst. However, by Day 3, the early and late isolates showed comparable levels of survival. By Day 7, the late isolate demonstrated significantly increased survival in U937 macrophages (p < 0.05), indicating a trend towards longer term survival inside macrophage cells. Unfortunately, P13 isolates were found to be resistant to 1 mg/mL of amikacin so macrophage uptake and survival could not be assessed using this method.

### Late infection isolates show changes in proteins associated with antibiotic resistance and intracellular survival

In order to further probe potential mechanisms for these changes in phenotype, whole proteome analysis was carried out on each pair of sequential isolates, comparing the proteome of the early isolate with that of the late isolate, to identify proteins which were significantly altered in abundance in late patient isolates which may contribute to these adaptations (**Table 3**). Due to the lack of a well-annotated proteome database for *M. abscessus*, *M. tuberculosis* strain H37Rv was used to assign functions.

**Table 3:**
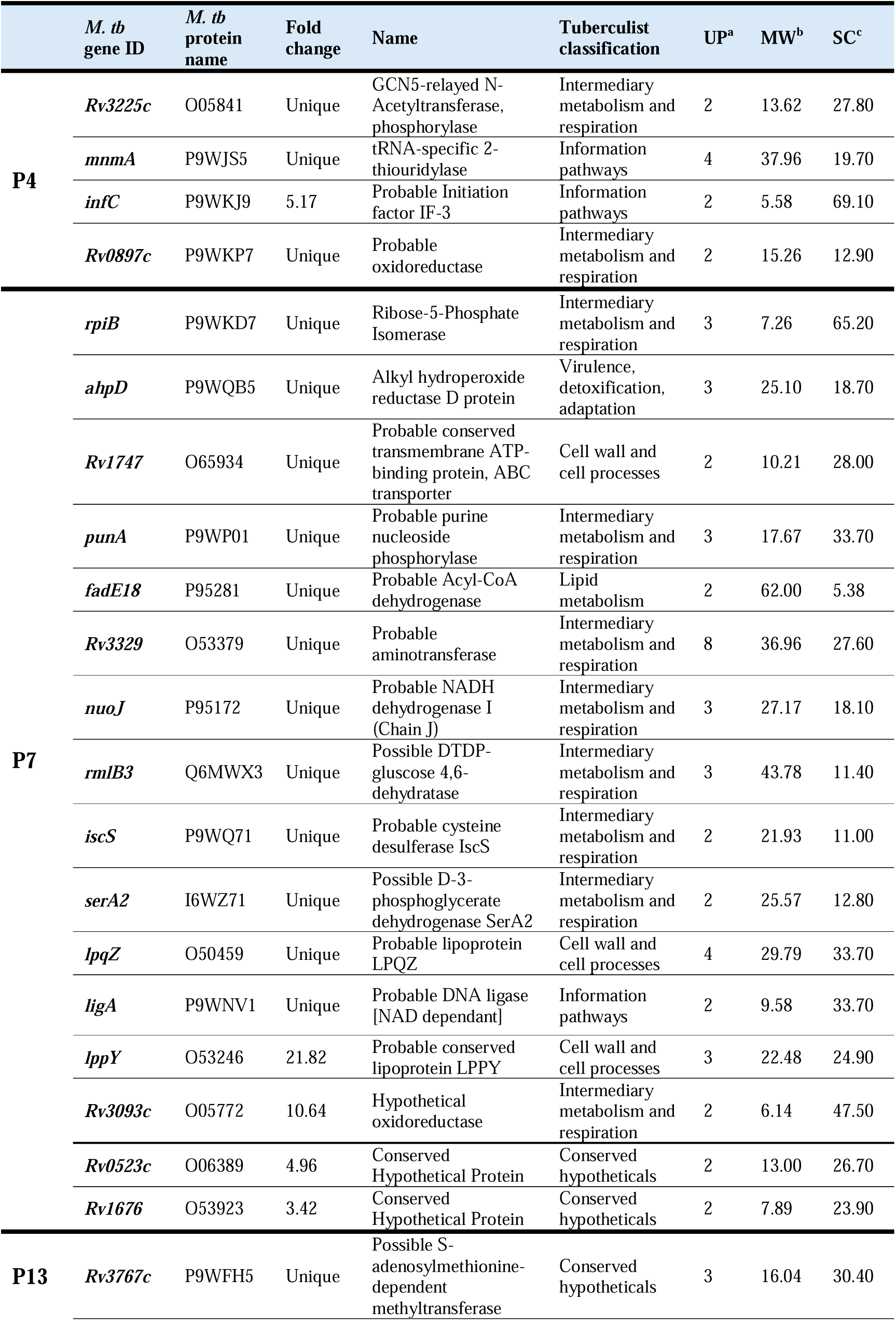
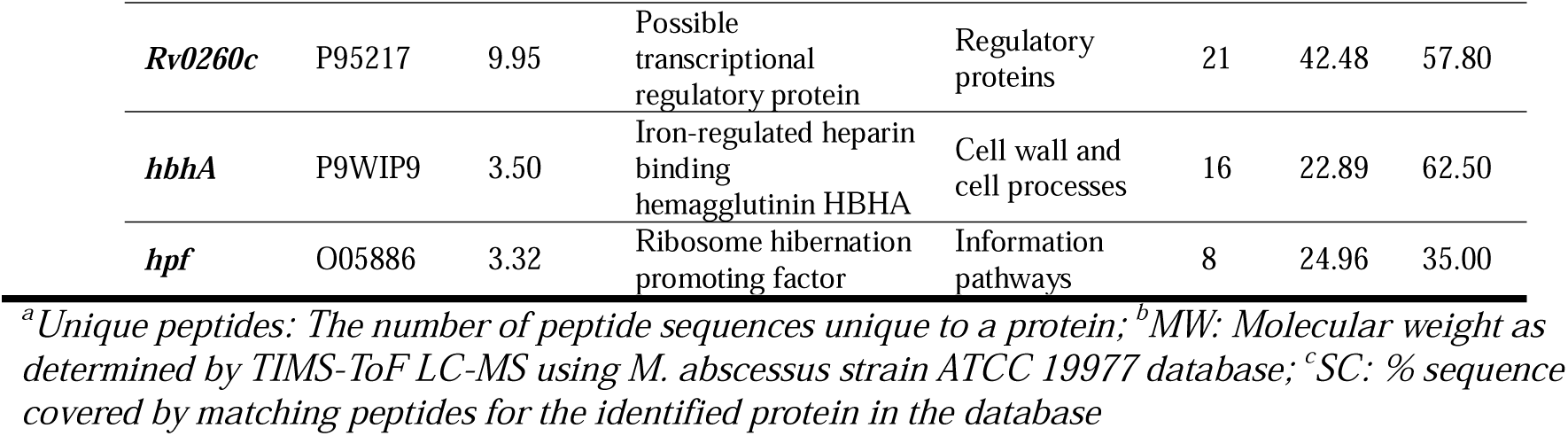
Proteins significantly increased in abundance by **≥** 1.5-fold in late infection isolates (*M.tb: M. tuberculosis*)

### P4

There appeared to be only 4 proteins that showed significant increase in abundance over the 85 day period between the early and late isolate in P4 (Table 3) and interestingly, three of these are reported to have roles in intracellular survival. Bifunctional GCN5-relayed N-acetyltransferase phosphorylase (AAC/APH) was unique to late isolate (undetectable in the early isolate) suggesting its expression was switched on over time of infection. This protein displays aminoglycoside kinase activity in *M. tuberculosis,* and is known to be involved in resistance to gentamicin, tobramycin and kanamycin (52, 53). tRNA-specific 2-thiouridylase MnmA was also unique to the late isolate. tRNA-specific 2-thiouridylases are important for correct codon recognition and stabilization of tRNA molecules (54). Deletion of *mnmA* attenuates growth of *M. tuberculosis* in macrophages, suggesting a role for this protein in facilitating intracellular growth in phagocytic cells (55). Probable Initiation factor IF-3 InfC was increased 5-fold in the late isolate. InfC has a known role in initiating protein translation and is induced during infection of human macrophage cells *in vitro* in *M. tuberculosis* (56, 57). Probable oxidoreductase, Rv0897c, was increased 4.5-fold in the late isolate. This hypothetical protein was previously found to accumulate in granulomatous lesions in TB (58).

The only protein showed significantly reduced abundance by >1.5-fold in the late P4 infection isolate, was a 2-enoylacyl coA hydratase (Ech), which was unique to the early P4 isolate and absent in the later isolate (Table S3). The enzyme is considered to play an important role in *M. tuberculosis* cells in lipid metabolism and the absence of this Ech suggests that gene expression was switched off in the late infection isolate. The *M. tuberculosis* genome contains approximately 21 Ech homologs that are expected to function as enoyl CoA hydratase/isomerases but the biochemical functions of many of these are not fully understood (59, 60).

### P7

Sixteen proteins were significantly increased in abundance in the P7 late isolate, including 12 that were undetectable in the early isolate (Table 3). This is notable as the time frame between early and late isolate was only 33 days. Ribose-5-Phosphate Isomerase (rpiB), an important enzyme in the pentose phosphate pathway in *M. tuberculosis*, was unique to late infection isolates (61). Alkyl hydroperoxide reductase D protein (AhpD) was also unique to the late isolate. AhpD is a critical element of the antioxidant defence system in *M. tuberculosis* as well as other pathogens (62). Together with AhpC, it provides a critical function particularly in isoniazid resistant isolates which tend to lack KatG, another catalase-peroxidase (63). As isoniazid is not typically prescribed for treatment of *M. abscessus* infection, it is thus probable that this increased abundance of *ahpD* is to provide extra antioxidant defence, potentially to resist the reactive oxygen species generated by many immune cells.

Unexpectedly, Rv3677, a beta lactamase was unique to the early infection isolate (Table S3). Another protein that was absent in the P7 late infection isolate was MprB, a two-component system sensor kinase that is a component of a cell envelope response network and interacts with the chaperone DnaK. In *M. smegmatis* deregulated production of MprB negatively regulates growth and interestingly MprAB activation is negatively regulated in the absence of stress (64). Its absence in the late infection P7 isolate may suggest that the niche this isolate was exposed to does not induce cell envelope stress, which is unexpected. However much more studies need to be performed on these proteins in *M. abscessus* and on its CF niche.

Eleven proteins were reduced in abundance in P7 isolates by 1.5-fold or more (Table S3). These include four components of the MCE lipid transporter (Mce2B, Mce3a, Mce3b and Mce1c) which were reduced in abundance by 1.77 to 5.15 fold. MCE (acronym for *mammalian cell entry*) proteins mediate lipid transport across the mycobacterial cell envelope and are considered important virulence factors, but may also other play roles apart from host-cell invasion(65). The Mce operon modifies *mce* gene expression in *M. tuberculosis* according to carbon source, and upon hypoxia, starvation, surface and oxidative stress (65).

### P13

Four proteins were significantly increased in abundance in the late P13 isolate, all of which have been reported to have a role in antibiotic resistance or host-cell interactions. The conserved hypothetical protein, Rv3767c, was uniquely detected in the late P13 isolate. Rv3767c is reportedly involved in resistance to pyrazinamide, a drug commonly used to treat TB (66). Possible transcriptional regulatory protein, Rv0260c was increased almost 10-fold.

Rv0260c interacts with the sensor kinase DosS of the DosRS system and is putatively involved in the response to stress (67). Iron-regulated heparin binding hemagglutinin HbhA, a known adhesin, was increased 3.5-fold in the late isolate. HbhA plays a role in the extrapulmonary dissemination of *M. tuberculosis* (68). Ribosome hibernation promoting factor hpf, showed a 3.3-fold increase in abundance in the late isolate. This protein is involved in the modulation of ribosome activity by binding to ribosomes and stabilising them to an inactive hibernating state in times of low nutrients or stress (69–71). Another ribosome hibernation promoting factor rafH is under the control of the DosR regulon and is upregulated in host macrophages in response to hypoxia and oxidative stress (71, 72). No proteins showed decreased abundance in the 13.6 isolate relative to the early infection isolates.

In order to identify if there were any potential common mechanisms of adaptation, we compared the proteins that showed increased abundance across the three sequential isolate pairs. This revealed that each isolate showed a unique proteomic profile to adapt to the specific pressures, which is reflective of the different environments of the individual patient CF lungs, which can show high degrees of heterogeneity even within the one patient(73). However, many proteins showing increased abundance had similar functions, suggesting that convergent processes may be used to respond to similar pressures (**Table 4**). In particular, adaptations in two processes were common across all the late infection isolates: proteins with roles in oxidative stress and intracellular survival; and proteins with roles in antibiotic and drug resistance. This suggests that these factors are critical for bacterial survival persistent infection. Lipid metabolism and cell wall biosynthesis were also observed in more than one late infection isolate.

**Table 4:** Comparisons of proteins significantly increased in abundance (≥ 1.5-fold) in late clinical isolates compared to their respective early isolates.

| Function/Pathway | P4 | P7 | P13 |
| --- | --- | --- | --- |
| Oxidative Stress/Intracellular survival | MnmA, Rv0897c | AhpD, IscS, Rv3093c | Rv0260c |
| Antibiotic/drug resistance | Rv3225c | Rv3066, | Rv3767c |
| Lipid metabolism |  | AccD2, FadE18 |  |
| Cell wall biosynthesis or component |  | RmlB3, Rv0302,<br>FabG2 | LpqH, MmpS3,<br>AcpM |

## Discussion

Pathoadaptation of *M. abscessus* is critical for its survival during chronic infection in the CF lung. Clinically relevant adaptations including reduced antibiotic susceptibility, increased attachment to host cells and increased intramacrophage survival were revealed in late infection isolates, demonstrating that the adaptation of *M. abscessus* over time of infection contributes to its persistence in the host and making it more difficult to eradicate (**Fig. 4**). The convergence in adaptations observed between isolates from three different CF patients suggests that intracellular survival and resistance to antibiotic pressure are key to surviving in the CF lung during chronic infection.

**Figure 4.**
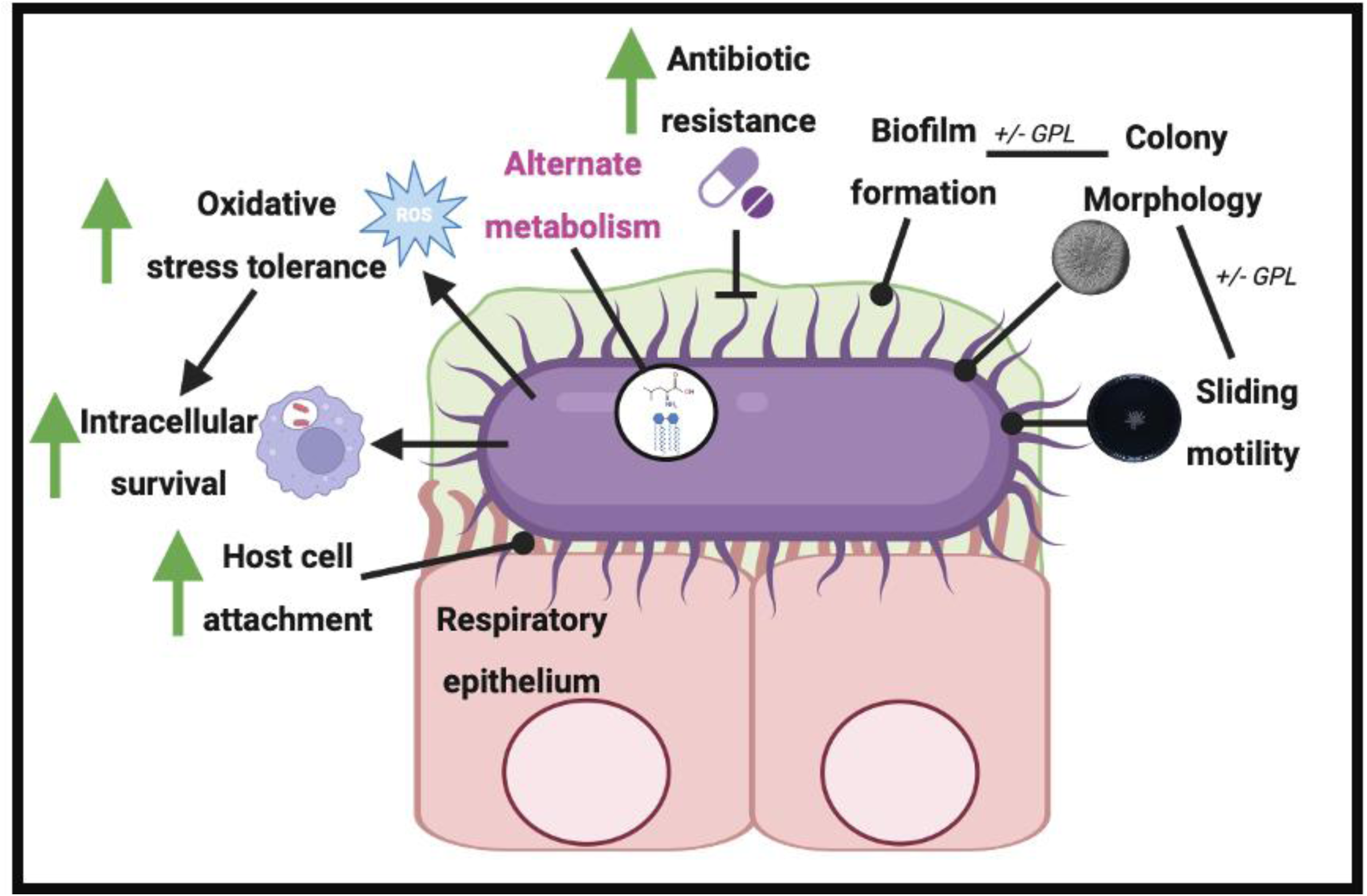
Phenotype and proteome adaptations demonstrated in late infection isolates. Changes marked with green arrows were shown by phenotypic assays and proteomic data while changes marked with pink text were identified from proteomic data only. Changes in biofilm formation, colony morphology and sliding motility were not seen in this data set but have been identified previously in clinical isolates. Created with Biorender.com.

Despite the difference in DCC origin, all three late isolates tested exhibited increased capacity to attach to CF epithelial cells relative to their respective early counterpart. We have previously shown that host cell attachment was also increased in two late *B. cenocepacia* sequential isolates from CF patients (48), suggesting that this might be a more general adaptation for these distinct yet challenging CF pathogens that share an intracellular lifestyle. Enhanced host cell attachment is likely to contribute to more effective colonisation of the CF lung and may be indicative of altered expression of surface proteins such as adhesins. Indeed, the enhanced abundance of iron-regulated heparin binding hemagglutinin in the late P13 isolate supports this theory. Moreover, the ability to survive intracellularly was also increased in late isolates from P4 and P7. The increased abundance of proteins associated with increased intracellular survival likely contributes to this adaptation. A limitation of this study is the lack of a well annotated proteome for *M. abscessus* relative to other CF pathogens thus there may be several other proteins that were altered in abundance and that were missed and the alterations in protein abundance observed are likely to underestimate the overall changes in proteome. This is an unfortunate ongoing challenge in the investigation of understudied bacterial pathogens, such as *M. abscessus*.

There are two interesting comparisons between isolate pairs which shed light on some of the paradigms of adaptation. Firstly, isolates from P7 demonstrated multiple significant changes over the course of just 33-days including: altered resistance to antibiotics; increased host cell attachment; increased survival in macrophages; and increased abundance of many proteins with diverse roles in infection and colonisation. This demonstrates that *M. abscessus* can respond relatively quickly to a new environmental niche. Secondly, it is apparent that the P13 early isolate was already adapted as there were very few changes observed over the time course between the early and late isolates; it appeared to be already highly resistant to antibiotics and showed a greater ability to adhere to CFBE cells than either of the other early isolates. It highlights that this isolate was perhaps already partially adapted to the CF lung niche at the time of acquisition. It should also be noted that the early isolate in each pair may not represent the initial or first isolate acquired by the patient. Although these isolates represent the earliest isolates since their transition to the adult CF unit for each of these patients, the patients may have acquired *M. abscessus* prior to transitioning from a paediatric facility and consequently the time of first acquisition is difficult to establish. It is possible that these isolates had already begun to adapt, which would have masked some of the changes observed. In the case of the P13 isolates, it is quite likely that the ‘early’ isolate was not the initial isolate acquired by the patient. Furthermore, while each isolate pair was confirmed to be clonal and potentially sequential via whole genome sequencing however, it is possible that multiple lineages may arise in the same patient. Isolation of colonising bacteria from sputum samples does not allow for sampling of specific areas of the lung and only one isolate per timepoint was selected, which likely does not represent the entire bacterial population.

The changes in proteome across the different late isolates reflect the different environments of the CF lung in individual patients, which show high degrees of heterogeneity between patients. Different CF mutations and modifier genes, co-infection with different populations of bacteria, different treatment options as well as a plethora of other environmental factors can infect disease severity in the CF lung and thus the pressures to which the colonising bacteria must adapt. This heterogeneity is reflected in the individual patient clinical data **(Table 1).** P4 and P7 are homozygous for the ΔF508 mutation and were prescribed modulator therapy in 2017/2018 which has contributed to their improved their lung function. In contrast, P13 is heterozygous for the c.1477C>T (p.Gln493X) mutation which is a functional Class I mutation (74) and consequently is not being treated with modulator and consequently P13 has experienced a continuous decline in FEV1 over time.

While these clinical data highlight the unique pressures which shape each CF patient lung environment, the convergence in phenotypes observed between isolates from three different patients, in addition to the increased abundance of proteins with roles in intracellular survival and antibiotic/resistance, stresses the importance of these adaptations for colonisation and survival in the CF lung environment.

The respiratory tract of people with CF or other underlying lung conditions is characterised by thickened mucus, chronic inflammation, and a constant bacterial burden which can lead to the development of hypoxic niches in the CF lung (8, 75, 76). It is likely that the natural habitat of *M. abscesses* in the environment may facilitate its survival in the CF lung as it must display exquisite adaptability to survive in the soil, water and in amoeba (77). These pre-adaptations may equip the isolates to better survive in the CF lung niche as well as intracellularly in macrophage cells (78).

In the last decade, CF modulator therapy has been transformative for people with CF with dramatically improved lung function and reduced exacerbations (79–81). However, recent studies have shown that while sputum density declines after the introduction of modulator therapy, most participants remain culture positive for CF pathogens after six months with some even becoming newly culture positive (79, 82). Pathogens were not fully eradicated and could continue to adapt albeit more slowly due a smaller population size (82, 83). This could be due to residual structural damage to the lung, failure to fully restore CFTR function and adaptation of these pathogens over time to stress to the point where they can survive in a ‘healthy’ CFTR-normalised lung (79). This highlights that adaptation during chronic infection remains a major issue for people with CF for which there are relatively few effective treatment options.

Treating pulmonary *M. abscessus* infections is highly challenging due to their resistance to aminoglycosides, rifamycins, tetracyclines, and β-lactams (84, 85). Consequently, managing *M. abscessus* infections often involves extended antimicrobial regimens lasting months to years (86). These prolonged treatments are associated with substantial side effects, including the risk of antibiotic toxicity and a high failure rate (87). Alternative treatments to target bacterial adaptation could have a significant synergistic effect with classical therapies and could allow these therapies to work as designed. Overall, these data show that *M. abscessus* adapts by enhancing its interactions with host cells and highlight that it has developed mechanisms to avoid clearance and facilitating persistence. Although these data increase our understanding of the adaptations that *M. abscessus* undergoes in the CF lung, additional studies are required to elucidate the mechanisms by which *M. abscessus* adapts to facilitate the development of novel therapeutics to combat this difficult to eradicate pathogen.

## 10. Author statements

### 10.1 Author contributions

Niamh Duggan: Investigation (lead); formal analysis (lead); methodology (lead); writing– original draft (lead); writing–review and editing (equal).

Joanna Drabinska: Formal analysis (supporting); investigation (equal); methodology (equal); writing–review and editing (supporting).

Ciarán J. Carey: Formal analysis (supporting); investigation (equal); methodology (equal); writing–review and editing (supporting Deirdre Keating: Formal analysis (supporting); investigation (supporting); methodology (supporting) Kirsten Schaffer: Resources (lead);supervision (supporting); writing–review and editing (supporting) Siobhán McClean: Conceptualization (lead); data curation (lead); funding acquisition (lead); investigation (equal); project administration (lead); resources (lead); supervision (lead); visualization (lead); writing–original draft (equal); writing–review and editing (lead).

## Ethical approval

N/A

## Consent for publication

N/A

