## Supplemental data for "From colonisation to chronicity: adaptation of *Mycobacterium abscessus* in the cystic fibrosis lung environment"

**Supplementary data**

**Table S1: Isolates selected for the construction of the phylogenetic tree and DCC clustering.** Taken from (24)

| **Sample** | **Subspecies** | **Cluster** | **Patient** | **Origin** | **Continent** | **Country** | **Location** | **Sample Date** | **Reference** |
| --- | --- | --- | --- | --- | --- | --- | --- | --- | --- |
| ERR3198402 | *M. a. abscessus* | DCC1 | GOSH18 | CF | Europe | UK | London_GOSH | 11/03/2008 | (88) |
| ERR4020109 | *M. a. abscessus* | DCC1 | Ireland_12 | CF | Europe | UK | Ireland_A | Not present | (89) |
| ERR459974 | *M. a. abscessus* | DCC1 | SVH_S | CF | Europe | UK | Dublin | Not present | (21) |
| ERR4020060 | *M. a. abscessus* | DCC2 | Ireland_11 | CF | Europe | UK | Ireland_E | Not present | (89) |
| ERR4020088 | *M. a. abscessus* | DCC2 | Ireland_14 | CF | Europe | UK | Ireland_A | Not present | (89) |
| ERR363330 | *M. a. abscessus* | DCC2 | AHL_C | CF | Europe | UK | Alder_Hey | 28/11/2011 | (21) |
| ERR4020112 | *M. a. abscessus* | DCC4 | Ireland_10 | CF | Europe | UK | Ireland_A | Not present | (89) |
| ERR340552 | *M. a. abscessus* | DCC4 | RBL_Q | CF | Europe | UK | London_Brompton | Not present | (21) |
| ERR459969 | *M. a. abscessus* | DCC4 | SVH_I | CF | Europe | UK | Dublin | Not present | (21) |
| SRR7800718 | *M. a. abscessus* | DCC5 | CF00039 | CF | North America | USA | TX | 27/06/2015 | (90) |
| ERR369139 | *M. a. abscessus* | DCC5 | DEN_Q | CF | Europe | Denmark | Copenhagen | Not present | (21) |
| ERR3198504 | *M. a. abscessus* | DCC5 | GOSH8 | CF | Europe | UK | London_GOSH | 29/06/2007 | (88) |
| ERR3198392 | *M. a. abscessus* | DCC5 | HVH43 | Non-CF | Europe | Spain | Barcelona_HVH | 23/02/2013 | (88) |
| ERR494976 | *M. a. abscessus* | DCC5 | SWE_B | CF | Europe | Sweden | Sweden | Not present | (21) |
| ERR363431 | *M. a. abscessus* | DCC5 | UNC_BK | CF | North America | USA | UNC_Chapel_Hill | 23/05/2008 | (21) |
| ERR373960 | *M. a. abscessus* | DCC5 | UNC_CU | CF | North America | USA | UNC_Chapel_Hill | 20/10/2005 | (21) |
| ERR373956 | *M. a. abscessus* | DCC5 | UNC_K | CF | North America | USA | UNC_Chapel_Hill | 07/07/2005 | (21) |

| Antimicrobial class | Antibiotic | Patient isolate | | | | | |
| --- | --- | --- | --- | --- | --- | --- | --- |
| ß-lactamases | Cephalosporins, penems, penams | 4.1 | 4.2 | 7.1 | 7.2 | 13.1 | 13.6 |
| Rifamycin |  | 4.1 | 4.2 | 7.1 | 7.2 | 13.1 | 13.6 |
| Glycopeptides |  | 4.1 | 4.2 | 7.1 | 7.2 | 13.1 | 13.6 |
| Aminoglycosides | Kanamycin | 4.1 | 4.2 | 7.1 | 7.2 |  | 13.6 |
| Aminoglycosides | Kanamycin A |  |  |  |  | 13.1 | 13.6 |
| Aminoglycosides | Neomycin |  |  |  |  |  | 13.6 (2 mutations) |
| Aminoglycosides | Gentamycin |  |  |  |  |  | 13.6 |
| Aminoglycosides | Gentamycin C |  |  |  |  |  | 13.6 |
| Aminoglycosides | Tobramycin |  |  |  |  |  | 13.6 (3 mutations) |
| Aminoglycosides | Amikacin |  |  |  |  |  | 13.6 (2 mutations) |
| Macrolides | Clarithromycin | 4.1 | 4.2 | 7.1 | 7.2 | 13.1 | 13.6 |
| Disinfecting agents/antiseptics |  |  |  | 7.2 | 7.2 |  |  |

**Table S2: Presence of genes associated with antimicrobial resistance.**

**Table S3: Proteins significantly decreased in abundance by ≥ 1.5-fold in late infection isolates**

|  |  |  | **Fold change** | **Name** | **Tuberculist classification** | **UP^a^** | **MW^b^** | **SC^c^** |
| --- | --- | --- | --- | --- | --- | --- | --- | --- |
| **P4** | **ChsH3** | Q6MWW2 | Unique | Possible 2-enoyl acyl-coA hydratase | Intermediary metabolism and respiration | 2 | 7.29 | 36.20 |
| **P7** | **FbiC** | P9WP77 | 1.55 | Probable F420 biosynthesis protein | Intermediary metabolism and respiration | 9 | 77.23 | 19.10 |
|  | **Rv2169c** | O53503 | 1.77 | Probable conserved transmembrane protein | Cell wall processes | 4 | 20.37 | 21.30 |
|  | **mce2B, Rv0590** | O07788 | 2.05 | Mce-family protein | Virulence, detoxification, adaptation | 2 | 38.26 | 7.90 |
|  | **cysNC, cysN** | P9WNM5 | 2.18 | Probable bifunctional enzyme cysn/cysc: sulfate adenyltransferase (subunit 1) + adenylylsulfate kinase | Intermediary metabolism and respiration | 5 | 18.24 | 48.60 |
|  | **recR, Rv3715c** | P9WHI3 | 2.34 | Probable recombination protein | Information pahways | 40.9 | 23.33 | 40.90 |
|  | **Mce3A,** | L7N698 | 2.40 | Mce-family protein mce3a | Virulence, detoxification, adaptation | 14 | 48.06 | 49.30 |
|  | **Rv1697** | O33198 | 2.88 | Conserved hypothetical protein | Conserved hypothetical | 8 | 39.92 | 30.00 |
|  | **Mce3B,** | O53968 | 3.33 | Mce-family protein mce3b | Virulence, detoxification, adaptation | 5 | 23.16 | 32.40 |
|  | **Rv3277** | P96882 | 4.19 | Probable conserved transmembrane protein | Cell wall processes | 2 | 27.04 | 8.70 |
|  | **YrbE4A,** | O53546 | 4.38 | Conserved hypothetical integral membrane protein yrbe4a. Possible ABC transporter. | Virulence, detoxification, adaptation | 4 | 30.05 | 13.00 |
|  | **Mce1C** | P9WP77 | 5.16 | Mce-family protein mce1c | Virulence, detoxification, adaptation | 5 | 38.12 | 22.10 |
|  | **MprB** | O53503 | Unique | Two component sensor kinase MprB | Regulatory proteins | 2 | 35.30 | 4.90 |
|  | **Rv3677c** | O07788 | Unique | Possible hydrolase | Intermediary metabolism and respiration | 5 | 27.52 | 36.00 |

*^a^Unique peptides: The number of peptide sequences unique to a protein; ^b^MW: Molecular weight as determined by TIMS-ToF LC-MS using M. abscessus* strain ATCC 19977 *database; ^C^SC: % sequence covered by matching peptides for the identified protein in the database*
